# Rapid Prototyping of Microfluidic Devices with Stereolithographic 3D Printing

**DOI:** 10.1101/2025.07.10.662041

**Authors:** Hunter G. Mason, Chih-Hsiang Hu, Leandro Soto Cordova, Ramin M. Hakami, Remi Veneziano

**Author notes:** H.G.M. and C.H. contributed equally.

## Abstract

3D printing has become a prevalent technology in many fields such as manufacturing, architecture, and electronics. This additive manufacturing technique is also widely used for biomedical research and clinical applications to prototype or assemble biomedical devices and tools. 3D printing-based strategies for biocompatible materials offer greater design flexibility, enhanced versatility, and faster results than traditional fabrication techniques, advantages that could be especially beneficial to the development of microfluidic chips. The ability to simply and efficiently produce new chip molds from computer aided design (CAD) models would significantly transform the development process and expand its accessibility by removing the need for more complex and expensive lithography methods. However, with standard processing strategies, the use of 3D printed molds for casting functioning chips is limited by the poor quality of prints achievable with widely available 3D printers. To mitigate this issue and facilitate rapid microfluidic device prototyping, we have developed a simple procedure to print microfluidic molds using a stereolithographic (SLA) printer and produce functional polydimethylsiloxane (PDMS) microfluidic chips with height and width feature dimensions as low as 75 µm. Molds printed using a commercially available liquid photopolymer-based resin and processed using our strategy exhibited high dimensional fidelity to intended designs and significantly reduced average surface roughness (< 3 µm). Here, we describe a streamlined post-print processing workflow for SLA molds and its efficacy in reducing surface roughness while preserving dimensional fidelity and then demonstrate its utility by prototyping and optimizing a microfluidic extracellular vesicle (EV)‐exchange platform.

**Graphical Abstract:** Rapid prototyping of microfluidic device features using stereolithographic 3D printing.

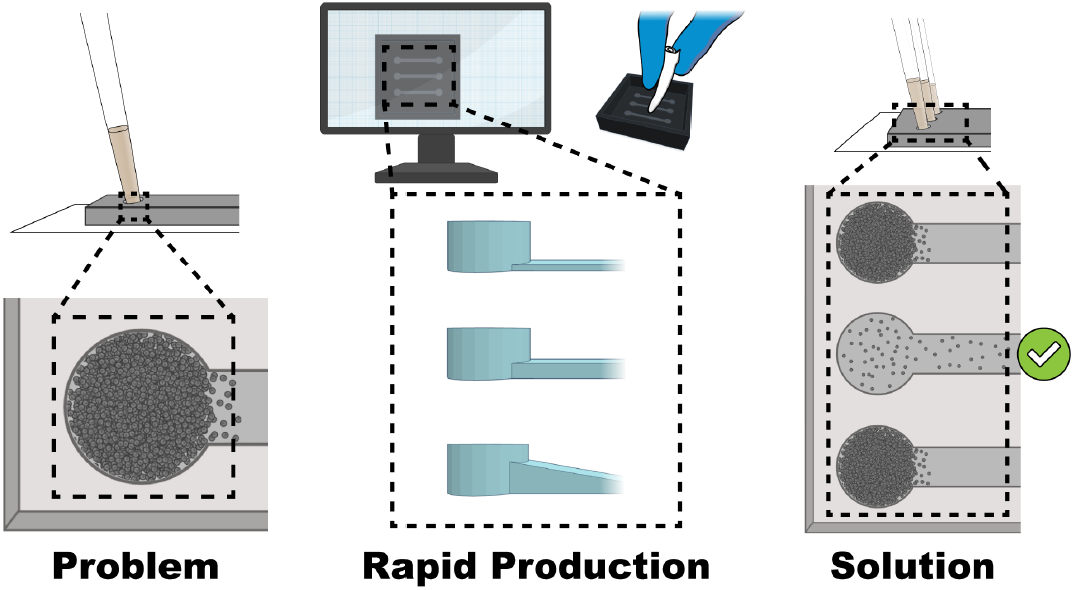

## 1. Introduction

Microfluidic devices have revolutionized biomedical research and particularly the way cell-based experiments are performed in several fields, including tissue engineering, biosensing, and drug discovery^1–4^. A single device can recapitulate several functions of an entire laboratory or of various biological systems^5–9^, offering simple high-throughput analysis in a finely controlled environment and with reduced sample volumes. These advantages not only reduce the cost of each assay but also facilitate multiplexing to enable screening or comparing an extensive array of compounds and conditions in parallel^10–12^. Microfluidic chips, often referred to as laboratory-on-chip (LOC), can be manufactured with different materials such as glass, silicon, ceramics, and polymers^13–16^. They are typically formed by the bonding of a molded base material with a flat surface (e.g., PDMS) onto another flat surface (e.g., glass or plastic) to create leak-proof, micro-thin grooves, channels, and/or small wells. A combination of interconnected microchannels, base materials properties, and precisely controlled sample infusion, achieve desired functional outputs such as guiding, separating, and mixing to fulfill specific experimental requirements.

Microfluidic devices used in biomedical research are prepared using a variety of technologies and methods such as photolithography, soft lithography, etching, hot embossing, and 3D printing^17,18^. The use of photolithography and soft lithography in tandem is the most prevalent fabrication strategy due to its high resolution and design flexibility. However, a major drawback is that not every laboratory has easy access to these costly technologies. In photolithography, ultra-flat material substrates such as silicon wafers are micropatterned by exposing spin-coated photoresist resins to UV light through photomasks. The resulting molds are then commonly used for soft lithography to imprint elastomeric materials such as PDMS. This master mold manufacturing process is costly and performed in clean room environments with specialized equipment, hindering the accessibility of these products to potential microfluidic chip users and limiting the capability to robustly prototype or iterate microfluidic chip designs. To circumvent these obstacles, cheaper and more accessible 3D printing strategies have been adopted to produce microfluidic chips and chip molds. As the irregularity and roughness of surfaces generated by standard 3D printers can interfere with important aspects of microfluidic devices including substrate bonding, fluid flow, and transparency, these strategies employ supplemental or alternative processing steps necessary to smoothen or transfer prints for use as functional microfluidic chips^19–22^. However, many of the simpler and efficacious processing strategies still involve multiple cumbersome steps and require additional equipment, or may rely on pre-coating the mold surfaces, thus affecting the precision and/or accuracy of the resulting feature dimensions^23–26^. Therefore, to provide a simple, rapid, and cost-effective solution for prototyping microfluidic devices, we developed and characterized a novel method, enabling the 3D printing of customizable microfluidic molds using an SLA printer. These master molds can be used to generate PDMS slabs with sufficiently smooth surfaces for permanent adhesion to glass substrates. Moreover, the processing is relatively non-destructive and precise, thus enabling high resolution printing to test critical features such as channel width and height.

Using our strategy, mold fabrication can be easily completed with virtually any commercially available SLA 3D printer and basic laboratory materials, and reagents. This enables the optimized procedure presented here to maintain the simplicity and low-cost of 3D printing while still mitigating the surface roughness of standard 3D printed devices. We demonstrate here the potential utility of our technique by rapidly prototyping and testing specific channel features to improve the design of one of our previously published microfluidic devices used for cell-cell communication studies^27^. Specifically, in our original design, cell clumping was often observed at the inlet-to-channel interface causing difficulties in controlling cell distribution and concentration throughout the full length of the channels. To test which feature adjustments could address this shortcoming, we applied our optimized processing procedure to quickly generate several chip molds with custom modifications in opening port angle and channel heights. Our studies showed that channel height was the main contributor to cell clogging and therefore we commissioned an improved SU-8 master mold of our chip designs with optimized channel heights without the need to order an intermediate prototyping mold, cutting the total effective cost in half in addition to rapidly achieving the desired outcome. The simplicity and robustness of our method should provide a universally available opportunity for a broader population of scientists to engage in microfluidic device-based research.

## 2 Results and discussion

### 2.1. Characterization of optimized processing of 3D printed molds and assembly of microfluidic chips

The use of 3D printed materials as molds for casting PDMS to be bonded to glass or other flat substrates is impeded by the resulting surface finishes. A standard procedure for processing fresh uncured prints typically involves repeated washes in an agitated solvent until the surface visually appears free of imperfections. However, this process leaves residual micro-sized resin clumps and other surface irregularities that are then transferred to the surfaces of molded PDMS slabs and can hinder PDMS-glass surface contact. To solve this problem, we developed a simple and robust surface smoothening method for SLA printed molds (Figure 1). Briefly, freshly printed molds are first wetted with 91% isopropyl alcohol (IPA) and gently wiped using a lint-free tissue. To limit the amount of force applied to fragile pre-cured channel structures, the tissue is pressed down on areas surrounding the channels for repeated wiping, allowing soft contact while limiting direct presses on channel surfaces that could damage delicate features. The molds are then dried by pressurized air and washed in a Form Wash with 91% IPA for 2 minutes. This process is performed three times to remove all residues on the surface of the molds before final drying step and UV curing in a Form Cure for 2 hours at 60 °C. The mold is initially placed with the channel side down for a few minutes before flipping it over to finish the curing process, which ensure thorough curing of the mold. By gently wiping prior to the final curing step, the surface is effectively sanded down without observable scratches that have been reported when sanding is done post-curing^25^. Each mold was designed as a square compartment with an inner 441 mm^2^ surface area available for the implemented chip designs (Figure S1-7) and 3 mm tall walls to retain the PDMS (Figure S8).

**Figure 1.**
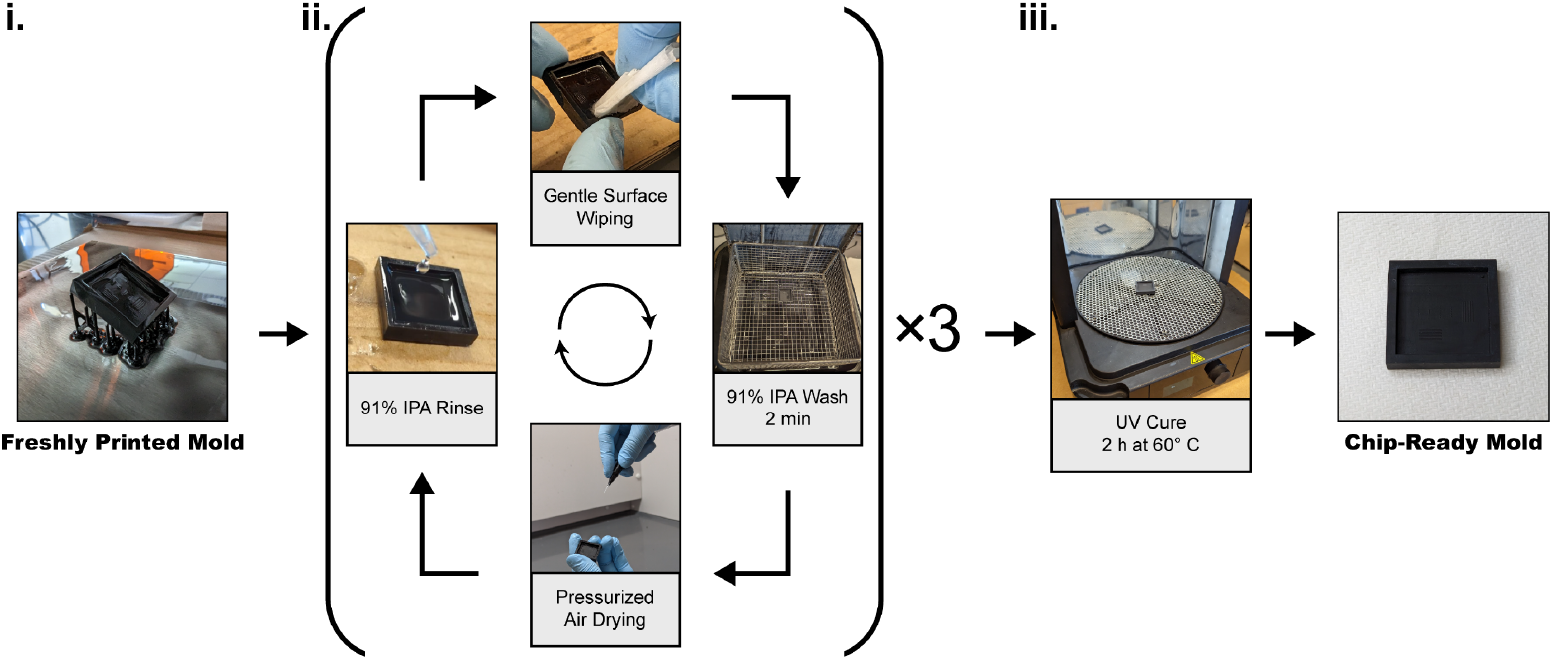
Optimized post-print processing workflow for 3D printed chip molds. Freshly printed chips (i) undergo 3 cycles of rinsing, surface wiping, washing, and drying (ii) followed by a final UV curing step (iii).

To evaluate the efficacy of our method, we produced two sets of 3D printed molds (Figure S1) that underwent standard post-print processing or our optimized processing and then analyzed the resulting molds and PDMS slabs. Standard processing was identical to the optimized procedure but without the wetting and gentle wiping of molds between washes in the Form Wash. First, PDMS chips fabricated using standard molds exhibited limited sealing to glass coverslips, causing notable fluid leakage (Figure 2a) or inadequate total bonding resulting in PDMS slabs peeling off with minimal force applied (Figure S9). In contrast, optimized PDMS chips are consistently and permanently sealed with only minor fluid leakage observed. Next, surface roughness was measured using 3D scanning laser microscopy. Topography images revealed that in the standard resin molds and PDMS slabs, there were several highly irregular surface mounds with ∼30-40 µm height ranges (Figure 2b). Comparatively, optimized surfaces displayed fewer notable height variations with reduced height ranges of ∼10-20 µm. Furthermore, analysis of the topography data showed a significant reduction in the average roughness (Sa), root-mean-square roughness (Sq), maximum peak height (Sp), and maximum valley depth (Sv) for optimized material surfaces (Figure 2c). Specifically, measured resin molds showed an average decrease in Sa and Sq of ∼57% and a decrease in Sv and Sp of ∼37% and ∼49%, respectively. These changes were also reflected downstream in the PDMS slabs, which showed an average decrease in Sa and Sq of ∼64% and a decrease in Sv and Sp of ∼60% and ∼29% respectively. Of note, the larger decrease in peak heights relative to valleys in the molds corresponded to a larger decrease in molded PDMS slab valley heights relative to peaks. Next, scanning electron microscopy (SEM) was used to visually compare the processed surfaces (Figure 2d). SEM imaging revealed the presence of large irregular clumps and mounds of resin in standard molds that were absent in the molds that underwent our optimized processing treatment. Similarly, the surfaces of standard PDMS slabs showed pits and crevices in the surface layer, especially surrounding channel imprints, that appeared diminished in size and quantity in optimized PDMS slabs. Furthermore, to determine if our processing technique was affecting the dimensional conformity of the print to the intended design, topography images of channels designed to be 500 µm x 500 µm and 275 µm x 500 µm (height and width respectively) were captured and analyzed (Figure S10). The average measured step height of the 500 µm and 275 µm channels were 517.6 ± 0.6 µm and 265.7 ± 2.3 µm respectively, both deviating ∼3.5% from the intended heights. The average widths of the 500 µm and 275 µm channels were approximately determined to be 535.8 ± 14.4 µm and 503.8 ± 7.054 µm respectively, deviating ∼7.1% and ∼0.8% from the intended width. These small variations are within the tolerance range for SLA printing and therefore should not affect the function of the microfluidic chips produced with these molds. Together, the topography and SEM image analyses suggest that the mold smoothening and enhanced PDMS-to-glass sealing are due to the removal of large, embedded resin clumps from the pre-cured surface that remain after standard processing, and that the removal of these clumps by gentle wiping does not significantly affect the final height and width of the print.

**Figure 2.**
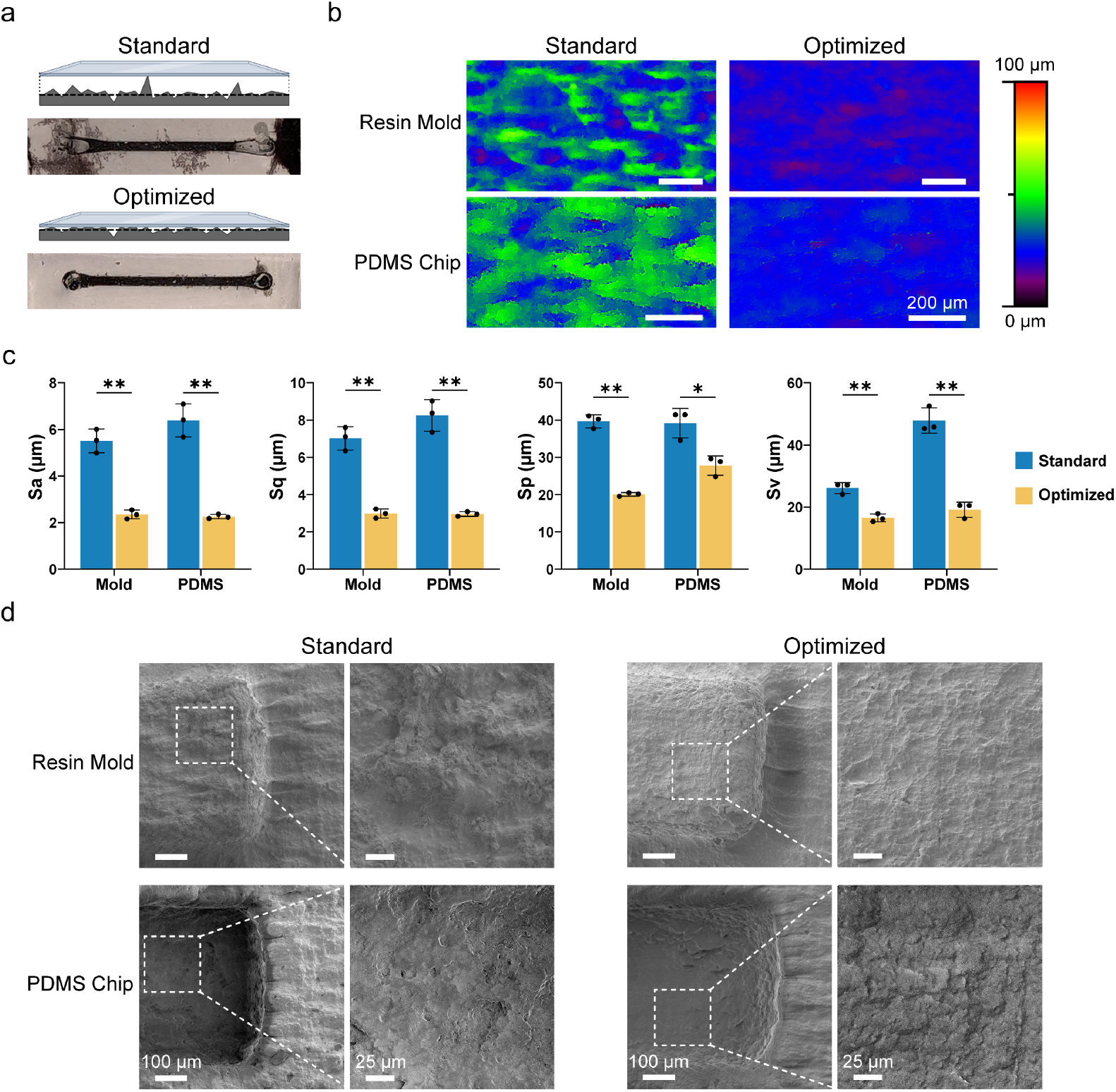
Surface characterization of molds produced by standard and optimized processing steps. (a) Glass bonding schematics and images of PDMS chips prepared using standard and optimized molds post-processing after black ink injection. (b) Laser scanning microscopy topography images of molds and PDMS slabs produced by standard and optimized procedures. (c) Comparison of average roughness (Sa), root mean square roughness (Sq), maximum valley depth (Sv), and maximum peak height (Sp) of the molds and PDMS slabs. ∼3 mm^2^ total area per replicate (n = 3) was scanned (Welch’s two-tailed *t*-test; error bars represent standard deviation of the mean; \**p* < 0.05; \*\**p* < 0.01). (d) SEM images of processed molds and gold coated PDMS slabs (Magnification: 140×; Inset Magnification 500×).

### 2.2. Rapid prototyping of features for microfluidic devices

The cost of developing a novel microfluidic chip system can potentially escalate by the need for feature and design testing due to the limited number of designs allotted per photomask plus the cost for materials and equipment time per new iteration. This cost is especially exacerbated when channel heights are assessed given the necessity of preliminary protocol optimization for fabricating photoresist regions at specified heights, the limited height range for different photoresists, and because producing molds with multiple heights on a single mask is cumbersome and greatly increases the time and cost invested during mold fabrication. Additionally, when in-house fabrication is not possible and mold fabrication needs to be commissioned at each design iteration, there can be a significant lag time in acquiring the molds for evaluation. Thus, we sought to demonstrate a real-world application of our optimized 3D printing strategy and its potential to facilitate efficient microfluidic chip prototyping. Cell accumulation in cylindrical microfluidic inlets has been previously observed with our microfluidic platform as well as others^28^. To alleviate this, we produced sets of 3D printed molds designed to assess various channel features and the resulting effects on cell distribution post-injection (Figure 3). The mold features tested include inlet opening angle (Figure S2), channel height (Figure S3), and sloped channel ceiling height (Figure S4), which is particularly complicated to achieve using classic photolithography techniques. During each test, suspensions of non-adherent U937 monocytes were gravity loaded into the channels for 1 hour followed by a 30-minute incubation period to allow the cells to settle in the channel. Cell distribution was then visually evaluated by brightfield microscopy. Three inlet opening types with a 311°, 241°, and 211° opening angle (Figure 3a) were first assessed wherein 311° represents standard port geometry and 241° and 211° correspond to expanded openings. Multiple repeat observations (n = 3) failed to show any substantial difference in post-injection cell distribution between the channel inlet opening variations. Next, four different channel heights were tested: 75 µm, 175 µm, 275 µm, and 375 µm (Figure 3b, Figure S11). Increased channel height was determined to be correlated with a reduction in port clogging, with channels 175 µm in height showing a noticeable reduction in port cell density and those 275 µm or 375 µm in height exhibiting no observable inlet accumulation. These results suggest that cell clogging is reduced as height increases until reaching a threshold between 175 µm and 275 µm. This is in accordance with the results of previous studies demonstrating that trapping and sedimentation of cells occurs at microfluidic inlet interfaces when channels sizes are less than 100-250 µm, depending on cell size and inlet design^29–32^ Lastly, slanted channels were designed and assessed to determine whether the height-based reduction in trapped cells is due to the enlarged openings at the inlet-channel interface or if the enhancement is dependent on the height of the full channel (Figure 3c). Two different step-wise slope designs with varying height gradations were tested. The results showed that injected cell accumulation was not reduced in either of these designs.

**Figure 3.**
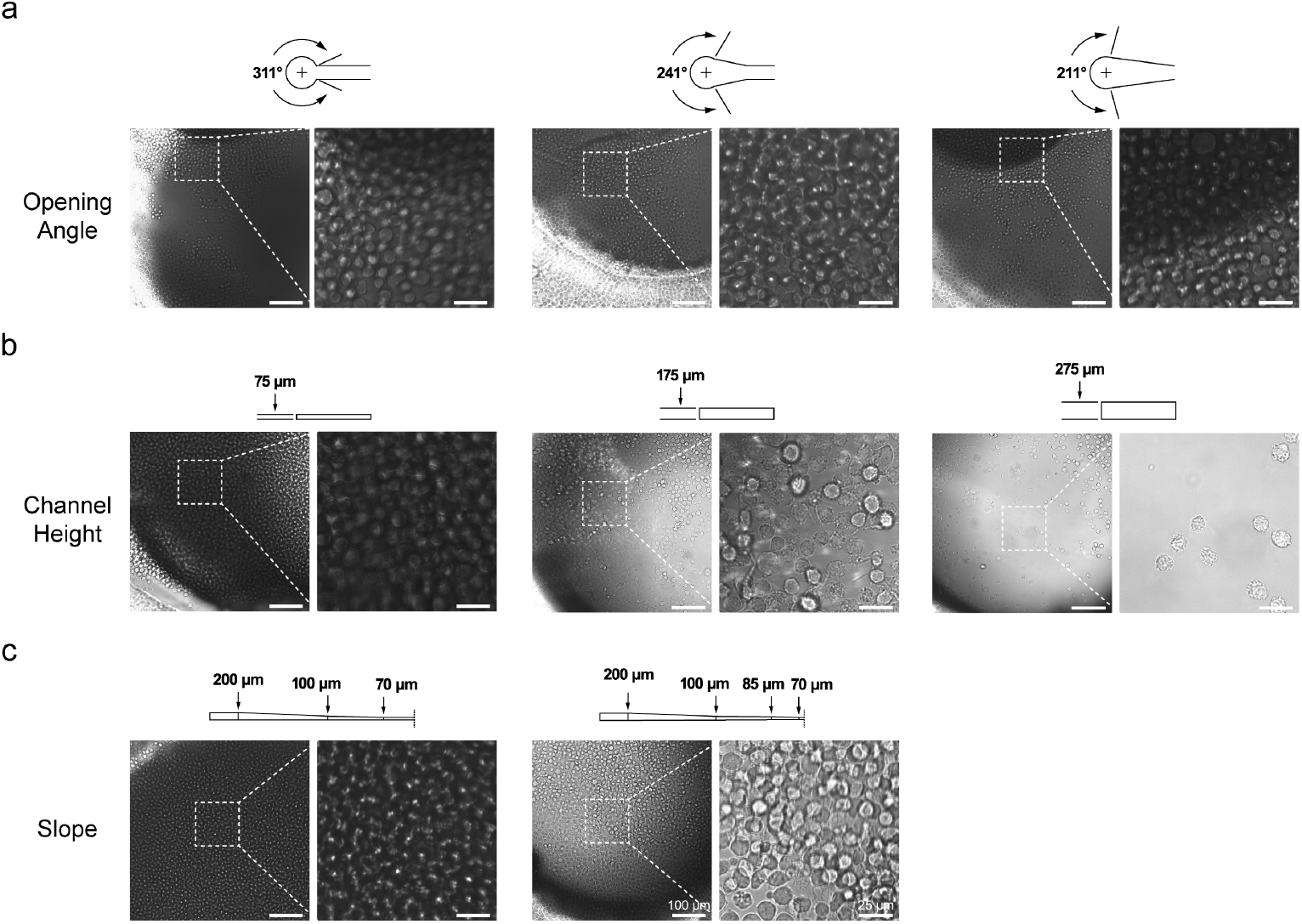
Rapid prototyping of chip features to assess effects on loaded cell distribution. U937 cells were gravity loaded for 1 h into the microfluidic device channel with varying (a) inlet port angles, (b) channel heights, and (c) sloping channel heights. Brightfield images of the channel inlets were taken 30 min post-loading to assess inlet cell accumulation (Magnification 600×).

Next, we applied our 3D printed prototype findings to design and fabricate an enhanced SU-8 photoresist mold. A replica design of our previously reported SU-8 molds^27^ was commissioned but with channel heights increased from 75 µm to 275 µm (Figure S5). Both the original and height-optimized molds were then used to fabricate PDMS chips and were visualized after gravity feeding the cells (Figure 4). In agreement with the prototype chip results, cell trapping observed at the inlets of the 75 µm chips was absent in the updated 275 µm chips. Thus, we have demonstrated that our 3D printing strategy can help mitigate the cost of refining microfluidic chip designs and streamline the development process towards achieving a final desirable product design.

**Figure 4.**
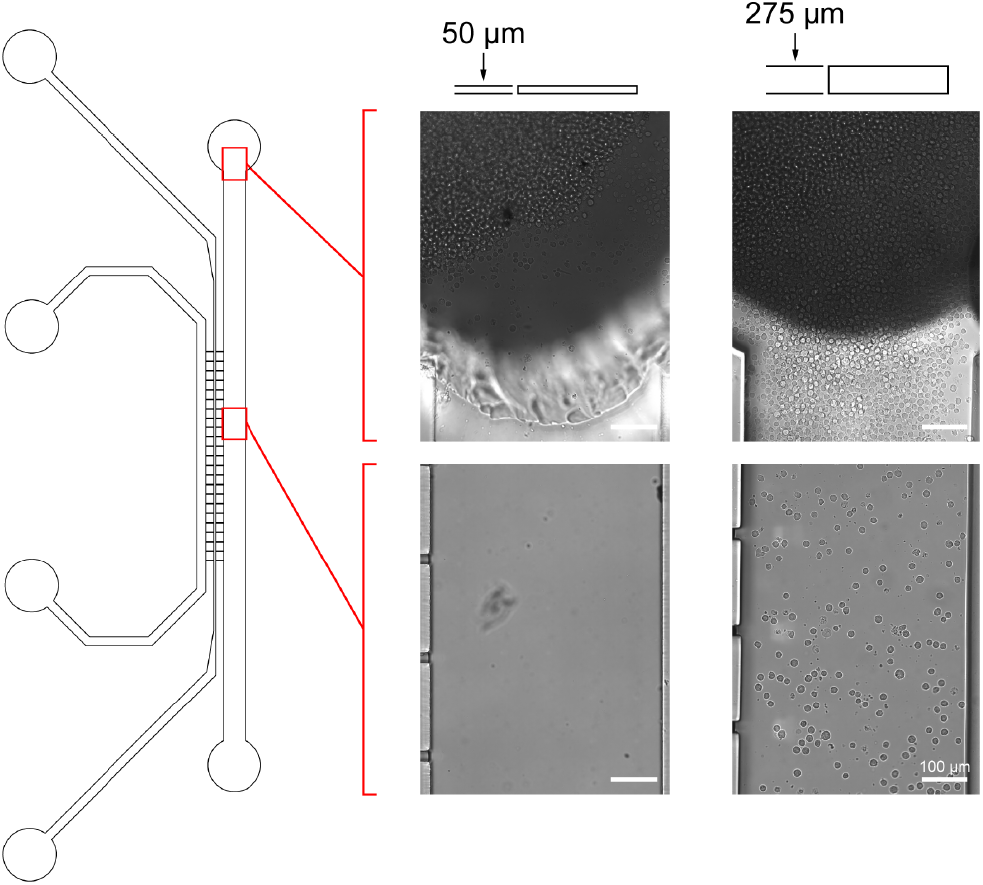
Application of 3D printed prototyping findings to SU-8 silicon wafer molds. Microfluidic chips were produced using the original 50 µm channel height mold and the newly fabricated 275 µm channel height SU-8 molds. U937 cells were gravity loaded for 1 hr and brightfield images of the channels were taken 30 min post-loading to visualize cell distribution (Magnification 600×).

### 2.3. Assessing the efficacy of our strategy with a biological application

One of the prevalent obstacles in the wider adoption and utilization of microfluidic chips by many research and teaching labs is the upfront cost investment and accessibility of equipment or facilities to prepare molds via photolithography. To verify the practicality and efficiency of our 3D printing strategy in transforming conceptual microfluidic chip designs into functioning systems, we used one of our previously published microfluidic chip models built onto SU-8 molded silicon wafer chips^27^. Briefly, the chip platform selected is set up to allow the monitoringof extracellular vesicle (EV) exchange between cells located in distinct channels towards deciphering regulatory functions of EVs in homeostasis^33^ and in a variety of pathological conditions^34^. In this microfluidic chip, the cell channels are connected by a fenestrated Matrigel-filled channel that selectively permits diffusion of particles based on hydrogel pore size and keeps the distinct cell populations separated. Due to the limitations of the 3D printer used in this study, as well as the choice we made to use standard resin, a 1-to-1 replica of our previous chip design (Figure S5) was not feasible. The 200 µm gap distance between the rib-like channels implemented in the original silicon wafer molds results in the cured resin of adjacent channels fusing together. Therefore, we first designed and produced a set of chip molds with incremental channel gaps, increasing the distance from 200 µm to 575 µm, to determine the minimum tolerated distancebetween adjacent printed channels (Figure S6). With a 3D laser scanning microscope, we determined that a 550 µm gap length was sufficient to facilitate printing distinct channels (Figure S12) and implemented it in the design of the final 3D printed EV exchange chip (Figure S7). This is consistent with the minimum recommended distance between adjacent channels for SLA printed molds. *In vitro* validation of the platform functionality was performed by loading U937 cells into the recipient channel of the chip followed by the injection of EVs labeled with CellMask Deep Red (CMDR) into the donor channel. The cells were then visualized at 4 hours post-EV injection and EV uptake in recipient cells was observed (Figure 5). Despite the design differences, the observed diffusion and uptake of EVs by recipient cells was functionally identical to that seen in our original chip platform (Figure 5). Thus, we have demonstrated that our optimized 3D printing processing technique can produce inexpensive but operational chips for validating the viability of conceptual chip designs. Lastly, while the printable resolution of channel features was constrained here by the available 3D printer, the use of higher fidelity printers with enhanced resolution and specialized resin may allow for prototyping exact replicates of final microfluidic chip designs when combined with our processing strategy.

**Figure 5.**
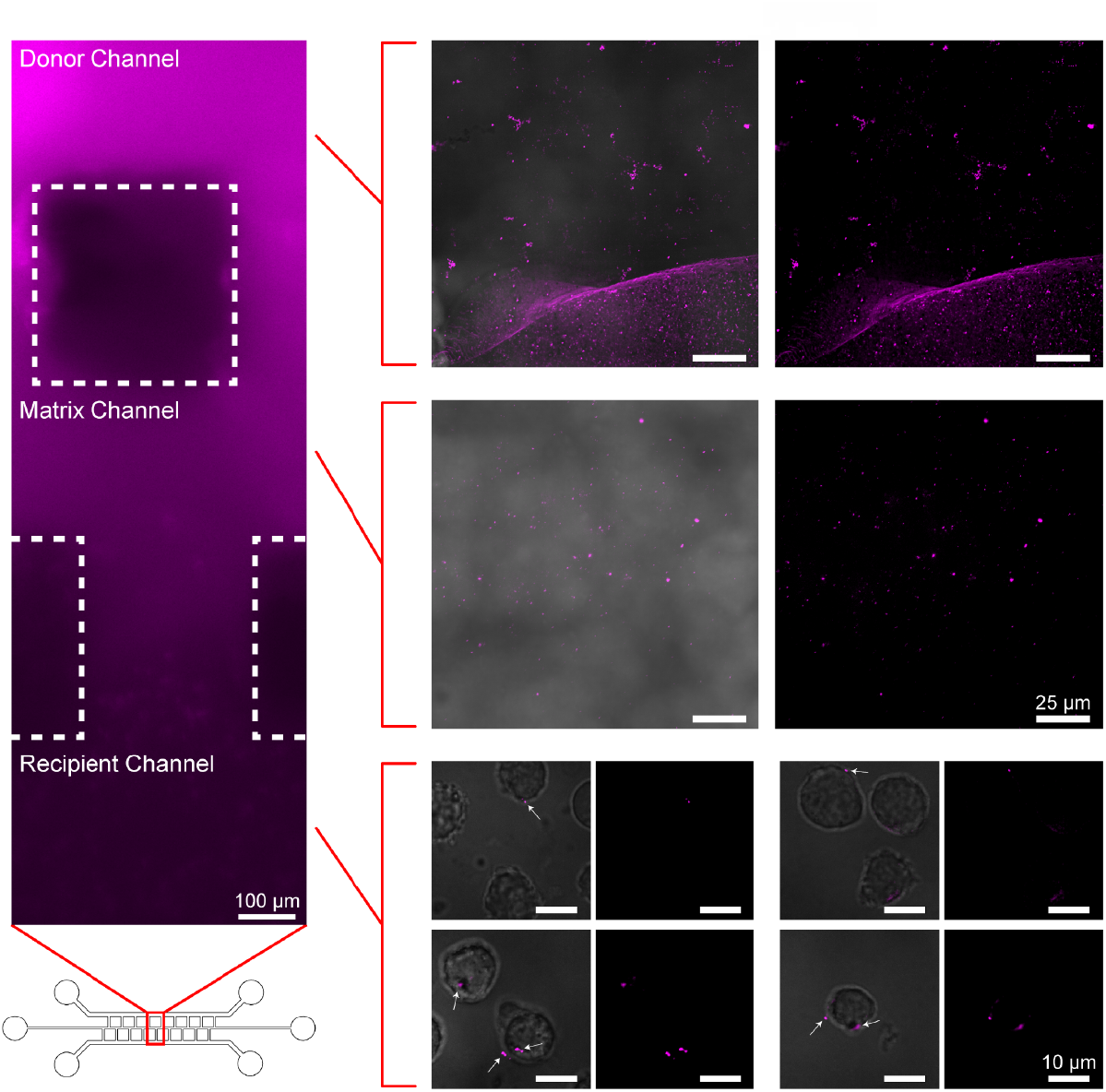
Demonstration of proof-of-concept 3D printing-based prototyping using an EV exchange microfluidic chip model. U937 cells were loaded in the recipient channel of the chip and subsequently CMDR-stained EVs were injected into the chip donor channel. The donor channel (top right), Matrigel-filled matrix channel (middle right), and recipient channel cells (bottom right) were visualized for the presence of EVs. EV-cell interactions and EV uptake into recipient U937 cells are indicated by white arrows. Brightfield and Cy5 emission filter images of the channels were taken 4 hours post EV injection (Left: Magnification 100×. Right: Magnification 1000×).

## 3. Conclusion

A novel strategy for accessible and facile production of microfluidic chip molds using SLA 3D printing was developed, characterized, and successfully tested. The optimized post-processing strategy developed can effectively smoothen the surface of printed materials, leading to ∼57% and ∼64% reduction in average surface roughness of molds and PDMS slabs respectively and without compromising the desired height or width of the print. We showed that this enhancement is sufficient to enable complete bonding of PDMS to glass with minimal fluid leakage to the periphery of molded channels. This processing strategy was validated in its utility by enabling swift and inexpensive production of multiple chip prototypesthat were used to resolve a previously observed cell congestion issue. The cell distribution results obtained using the 3D printed mold prototypes were verified by experimental evaluation of a newly fabricated SU-8 mold that incorporated our findings. Furthermore, the ability of this strategy to facilitate the development of novel functional chip platforms was established by the reproduction of an EV exchange platform. Thus, we have demonstrated its potential as an alternative to photolithography-based mold prototyping, with significantly reduced cost and time to analysis. This strategy should provide research teams without access to photolithography equipment and/or adequate funding a means of trialing novel microfluidic chip systems or prototyping design features to enhance the performance of existing platforms. Moreover, the simplicity and low cost of processing and printing of the molds could allow the introduction of custom microfluidic chips into a lab classroom as a hands-on teaching tool.

## 4. Methods

### 4.1. Mold design and Printing

The microfluidic chip molds were designed in AutoCAD Fusion 360 and exported as STL files. For the sloped height channels, a sequence of points with gradually varied heights were defined and connected. The files were uploaded to PreForm, a 3D slicer software for FormLabs, and each mold design was slanted approximately 45 ° with support added to the back side of the mold and channels facing upwards. Default settings were used with layer thickness set to 0.025 mm for maximized accuracy and Black V4 as selected resin type for its versatility. The resulting FORM files were uploaded to a Form 3+ Low Force Stereolithography (LFS) printer loaded with FormLabs Black V4 resin and printed at room temperature.

### 4.2. Removing and Smoothing Process

Freshly printed molds were removed from the Build Platform 2 by carefully bending the flexible platform and then underwent either standard or optimized processing as described in our results (Figure 1). All processing steps besides UV curing were done at room temperature. The support was removed prior to the treatment process. A Form Wash with 91% IPA and Form Cure were used for agitated solvent washes and UV curing respectively.

### 4.3. PDMS Chip Production

Before PDMS casting, processed molds were first cleaned by rinsing with 70% ethanol followed by drying using compressed air. To ensure that residual ethanol was fully dried and to alleviate reported inhibition of PDMS curing by uncured SLA resins^26,35^, the molds were then baked for 1 h at 70 °C. Polydimethylsiloxane (PDMS; Sylgard 184.) (Electron Microscopy Sciences, 24236-10) was harshly stirred at a 10:1 (m/m) ratio of base to curing agent and poured into the molds. The molds were subsequently placed into a vacuum chamber for 30 min, or until the PDMS was fully degassed, and then baked at 70 °C for 1 h. A scalpel was used to cut the PDMS slabs from the molds and then each cell channel inlet and outlet port was punched using a 1.2 mm sample punch (Ted Pella, 15115-4). To remove excess PDMS debris, the chips were then rinsed with 70% ethanol, ultrasonicated in ultrapure water for 5 min, and dried using compressed air. The chip surfaces and glass coverslips (VWR, 48393-081) were plasma activated for ∼10 seconds at max voltage settings using a handheld corona treater (Electro-Technic Products, BD-20AC), pressed firmly together ensuring no air gaps, and baked at 70 °C for 1 h for bonding. For EV exchange experiments, 3 μl of Matrigel (Corning, 354230) was injected into the middle channel and allowed to polymerize for 30 min at 37 °C. After polymerization, the cell channels were subsequently rinsed and filled with PBS.

### 4.4. 3D Scanning Laser Microscopy

Molds and PDMS slabs were measured using an Olympus LEXT OSL4100. Prior to measuring, PDMS slabs were gold sputter coated at 25 mA for 30 s using a Denton Desk V due to the transparency of PDMS. Surface roughness measurements were taken using an Olympus LMPLFLN50 (50× objective) lens. Three 1 mm^2^ regions were randomly chosen and scanned per mold for three different molds. All other measurements were performed using an Olympus MPLFLN10 (10× objective) lens. Scans were stitched, tilt-adjusted, and analyzed using the included OSL4100 software.

### 4.5. Scanning Electron Microscopy (SEM)

Molds and PDMS slabs were mounted to aluminum stubs (RaveScientific, RS-MN-10-005112-50) using adhesive carbon tabs (Electron Microscopy Sciences, 24236-10). Samples were then gold sputter coated at 25 mA for 30 s using a Denton Desk V. Images were captured with a JEOL JSM-7200F at an accelerating voltage range of 2 kV.

### 4.6. Cell Maintenance and Chip Loading

U937 cells (kind gift of Dr. Michael S. Strano’s laboratory at MIT) were maintained inside a tissue culture incubator (37 °C, 5% CO2) in RPMI 1640 medium with L-glutamine and HEPES (Corning, 10-041-CV) and supplemented with 10% heat-inactivated fetal bovine serum (FBS) (Corning, 35-010-CV). Vero cells (ATCC, CCL-81) were maintained inside a tissue culture incubator (37 °C, 5% CO2) in Dulbecco’s modified minimum essential medium (DMEM) with HEPES, high glucose, and L-glutamine (Corning, 10-027-CV). Before loading, cell count measurements were performed using a Luna Automated Cell Counter (Logos Biosystems, L10001) in fluorescence mode after mixing the cells with AO/PI dye (VitaScientific, F23001). For cell loading into chips, U937 cells were gravity fed into each cell channel; 40 μl of 1 × 10^6^ cells/ml were taken up into a 200 μl pipette tip which was then inserted into a cell channel inlet, ejected, and left in an upright position for 1 h under normal cell incubation conditions. The pipette tips were then gently removed. Loaded chips were placed in a tissue culture incubator (37 °C, 5% CO2) until use; cells in test chips for analyzing cell distribution were allowed to settle for 30 min prior to imaging, and EV recipient cells were incubated for 4 h prior to imaging.

### 4.7. Extracellular Vesicle Purification, Labeling, and Injection

For EV purification, we modified our previously published procedure^36^ using a slightly modified version of the protocol described by Tkach et al.^37^, and subsequently characterized the purified EVs by various marker analysis, electron microscopy (EM) studies, and nanoparticle tracking analysis (ZetaView) as previously described^36^. Briefly, Vero cells were expanded into T-300 flasks with DMEM supplemented with 10% EV-depleted FBS. At 48 h, the conditioned culture media was collected and spun down at 2000× g for 20 min at 4 °C. The resulting supernatant was then transferred and centrifuged again at 10,000× g for 30 min at 4 °C. The supernatant from the 10,000× g spin was filter-sterilized using a 0.22 μm bottle-top filter (Santa Cruz Biotechnology, sc-258381) and then concentrated to ∼1 ml using a 100 kDa MWCO Centricon Plus filter (Millipore Sigma, UFC710008). The concentrate was then loaded onto a 35 nm qEV1 column (Izon, IC1-35). Fraction volumes of 500 μl PBS were repeatedly loaded onto the columns, and fractions 9, 10, and 11 that contained EVs were collected and combined. The purified EVs were then labeled with 2.5 ug/ml CellMask Deep Red plasma membrane stain (ThermoFisher Scientific, C10046). Labeled EVs were then overlayed on a step-wise sucrose (VWR, 0335-1KG) density gradient prepared as described^36^, and the gradient was centrifuged at 274,000× g for 3h at 4 °C. The 1.151 g/ml and 1.118 g/ml fractions were then collected, subjected to 3 PBS buffer exchanges using a 100 kDa MWCO Amicon Ultra filter (Millipore Sigma, UFC910008), and were concentrated to a final volume of ∼200 μl. The purified EVs were then sterilized using a 0.20 μm sterile syringe filter (VWR, 28145-477) and used for EV exchange experiments within 30 min after purification. A 5 μl volume of the labeled EVs was pipette injected into the donor channel of the EV exchange chips after cell loading of the recipient channel and were incubated in the chips for 4 h before imaging.

### 4.8. Brightfield and Fluorescent Microscopy

Images of live cells and fluorescent EVs within the chips were acquired using a Nikon Ti2 microscope. Brightfield images of chips used in cell clogging tests were obtained with a 60× objective lens. EV exchange chips were imaged using brightfield and Cy5 filters, with a 10× objective lens and a 100× oil immersion objective lens.

### 4.9. Statistical Analysis

Mean, standard deviation, and statistical significance of values were determined and plotted using GraphPad Prism 10. Statistical significance was calculated by multiple Welch’s two-tailed *t*-test comparing samples that underwent standard or optimized processing.

## Author Contributions

^†^H.G.M. and C.H. contributed equally.

