## Supplementary Material for "Rapid Prototyping of Microfluidic Devices with Stereolithographic 3D Printing"

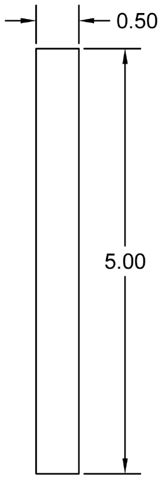


**Figure S1.** CAD drawing of the print used to compare the surface roughness and channel width/height produced by the standard and optimized processing strategies.

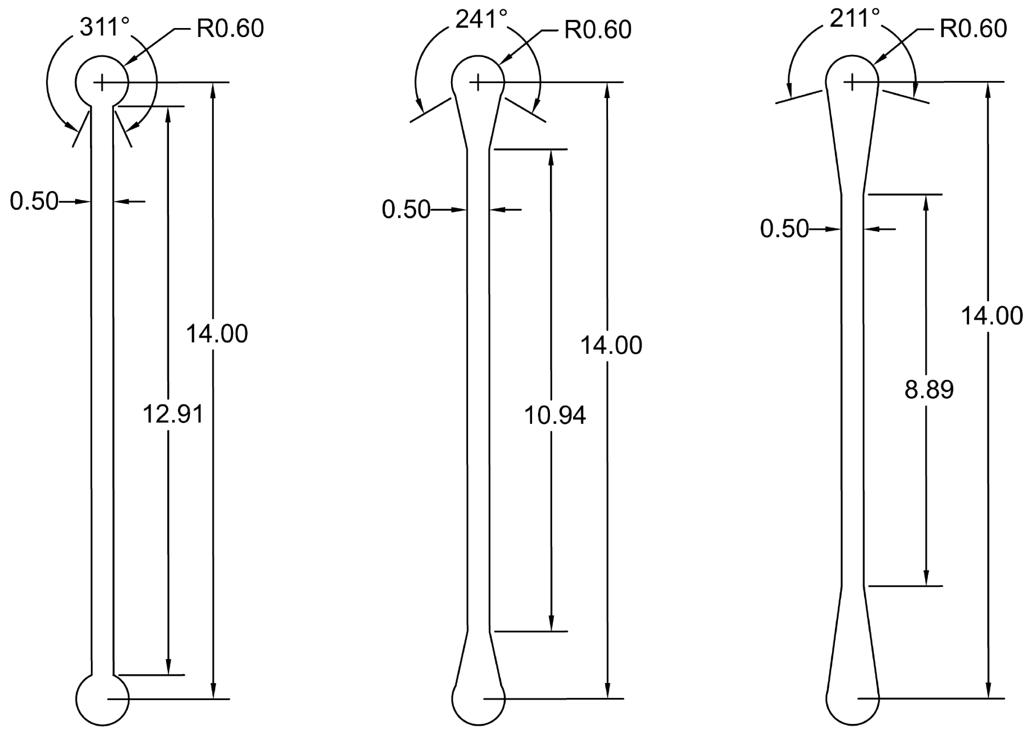


**Figure S2.** CAD drawings of the channels tested for analyzing the effect of port opening angles on cell distribution. Printed channel heights were set to 75 μm.

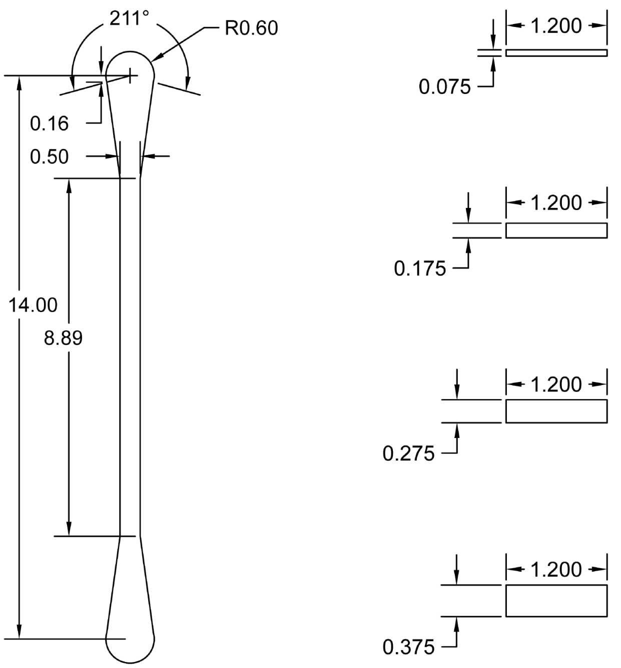


**Figure S3.** CAD drawings of the channels tested for analyzing the effect of channel heights on cell distribution.

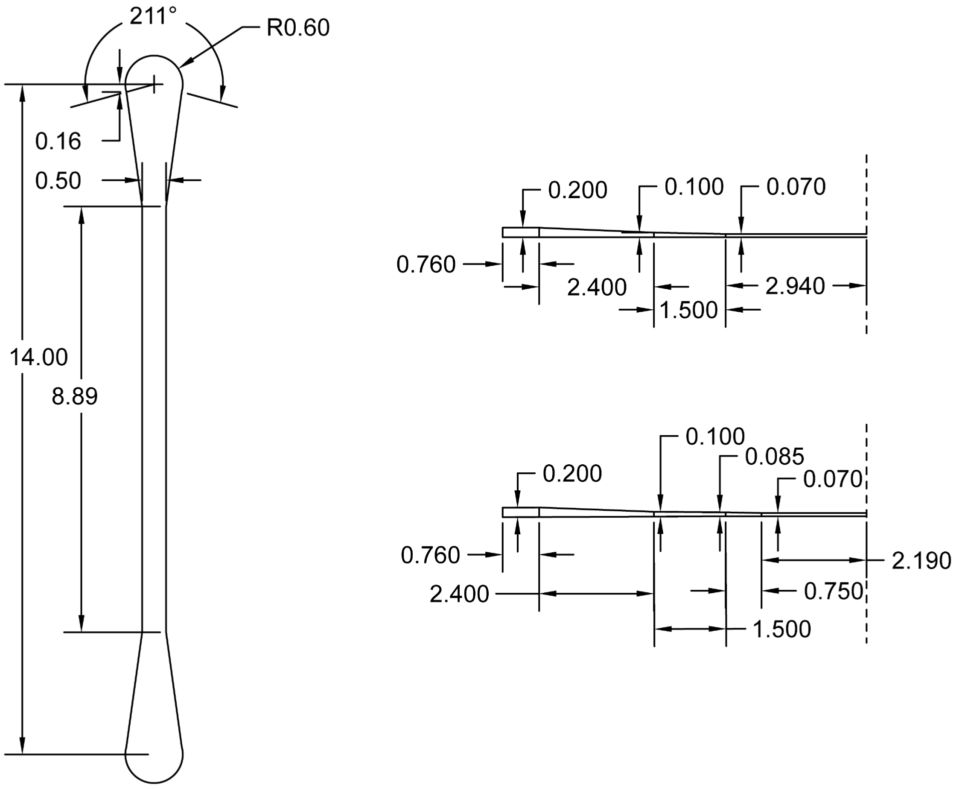


**Figure S4.** CAD drawings of the channels tested for analyzing the effect of sloped channel heights on cell distribution. The channels are mirrored across the dotted lines indicated in the height maps (right).

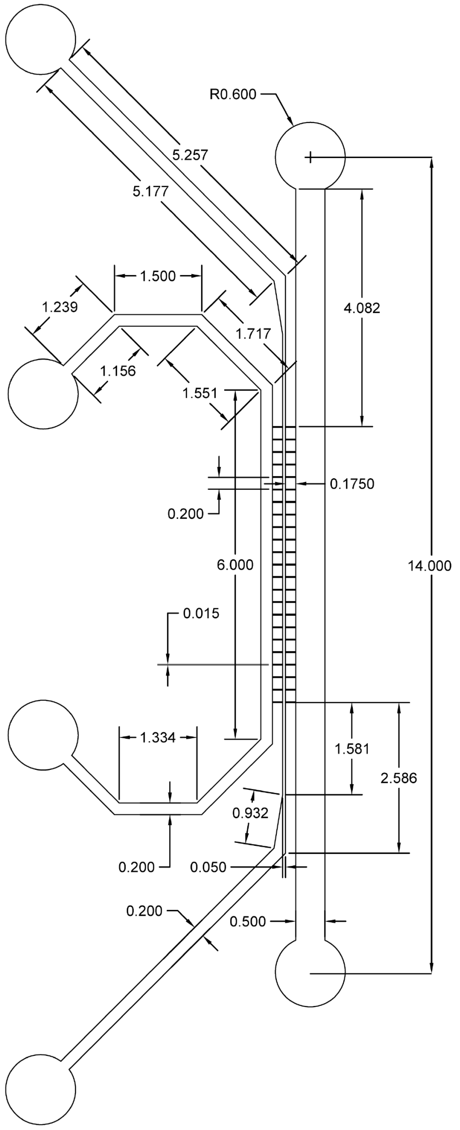


**Figure S5.** CAD drawing of SU-8 photoresist-based co-culture chips used for observation of live EV exchange between distinct cell channels. The original printed channel heights were 75 µm and the newly fabricated chip channel heights were 275 µm.

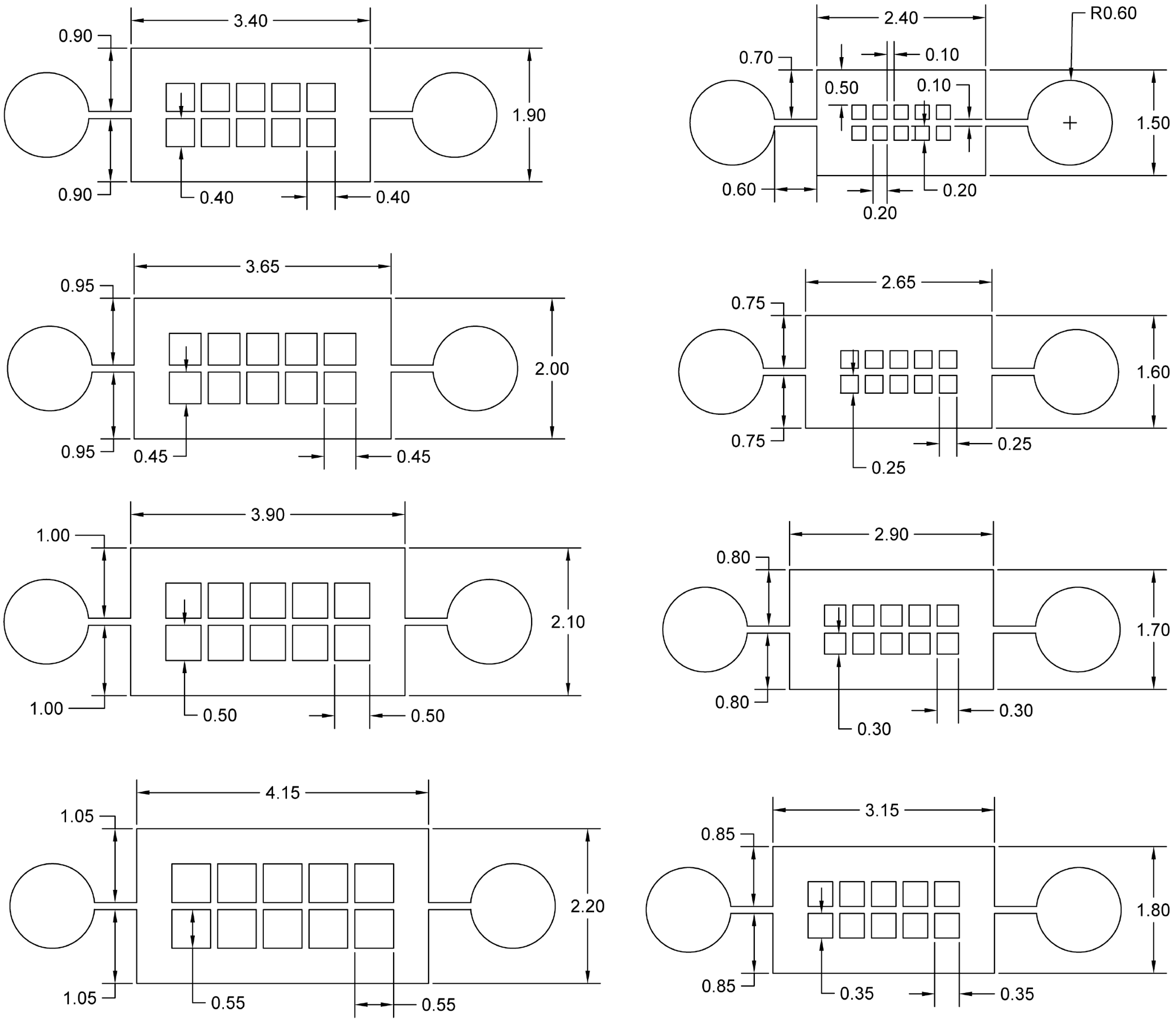


**Figure S6.** CAD drawings of the channels printed to evaluate the minimal printable resolution of chips tested for analyzing the effect of port opening angles on cell distribution. Printed channel heights were 275 µm.

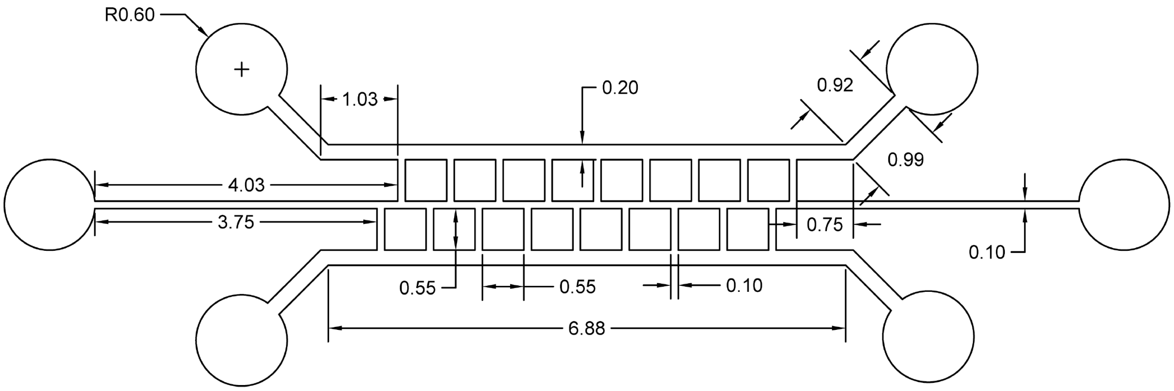


**Figure S7.** CAD drawing of the 3D printed EV exchange chip. The design is an altered version of the CAD drawing in Figure S5 that implements the minimum 3D printable resolution for channels. Printed channel heights were 275 μm.

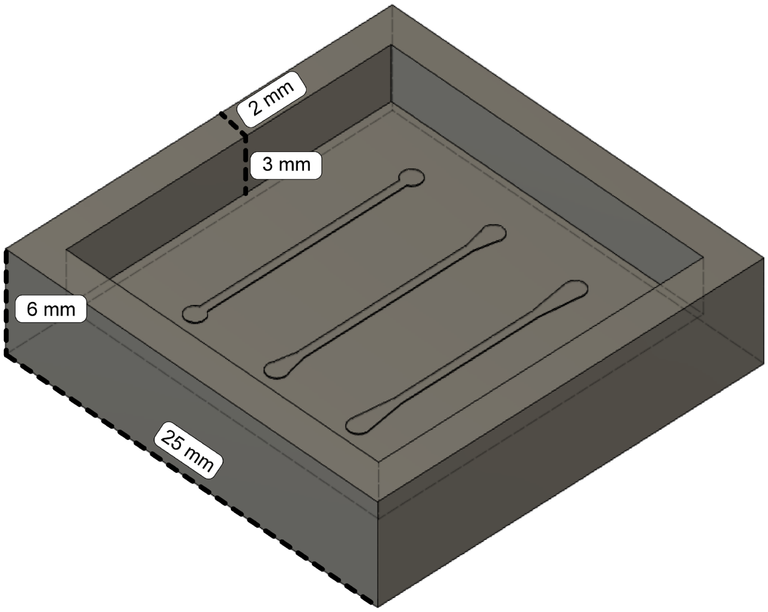


**Figure S8.** 3D model of a printed chip. PDMS is poured within the walls to produce ~2.5-3 mm thick chips.

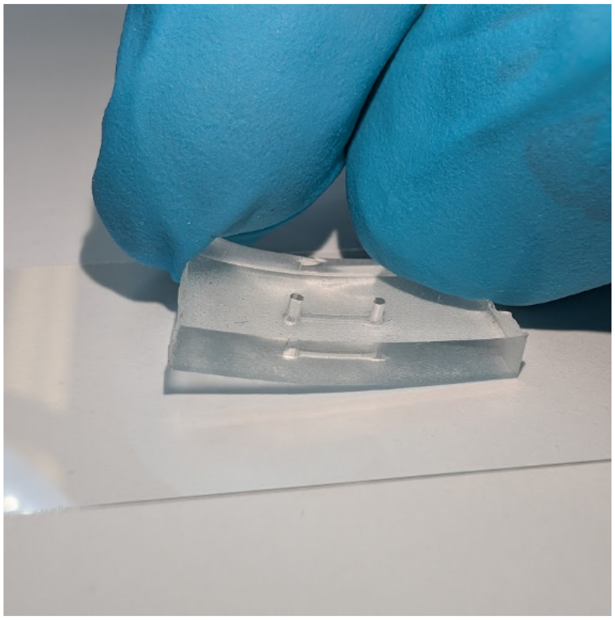

**Figure S9.** PDMS slab molded with standard processing easily lifts off the glass coverslip with minimal force applied after failing to covalently bond to glass due to molded PDMS surface properties.

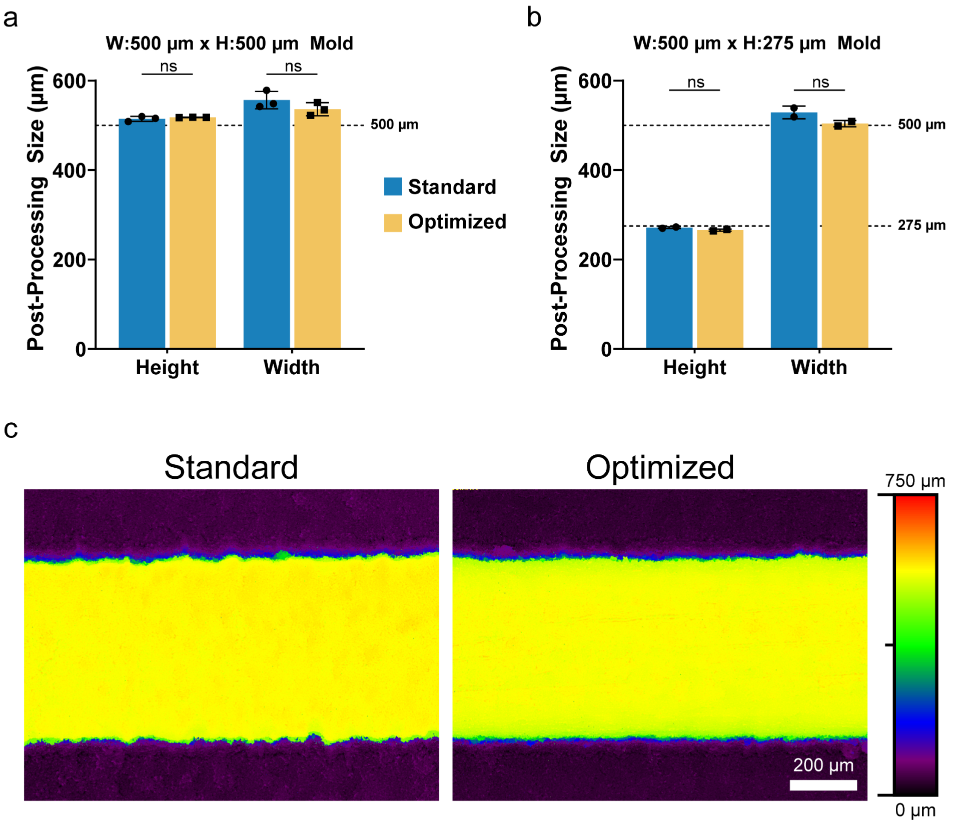


**Figure S10.** Comparison of channel step height and width between molds processed by standard and optimized steps. Step heights and widths were scanned and measured for (a) 500 µm height (n = 3) and (b) 275 µm height (n = 2) channel molds (Welch's two-tailed *t*-test; error bars represent standard deviation of the mean; ns denotes not significant). (c) Laser scanning microscopy topography images of 500 µm channel height molds produced by standard and optimized procedures.

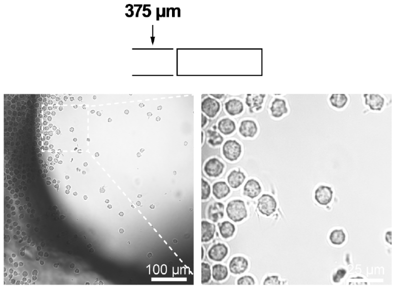


**Figure S11.** Assessment of cell distribution in the channel with 375 µm height. U937 cells were gravity loaded for 1 h and brightfield images of the channel inlets were taken 30 min post-loading to assess cell clogging (Magnification 600×).

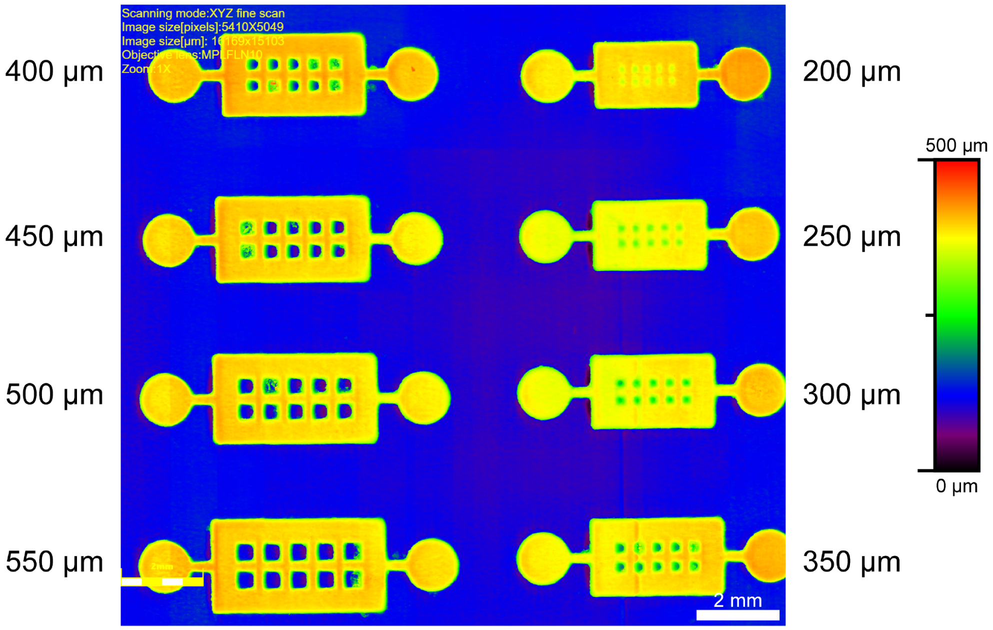


**Figure S12.** 3D printable channel resolution evaluation. Laser scanning microscopy topography image of the test mold after optimized processing is shown.
